# How to determine microbial lag phase duration?

**DOI:** 10.1101/2022.11.16.516631

**Authors:** Monika Opalek, Bogna J. Smug, Dominika Wloch-Salamon

## Abstract

The lag phase is a temporary non-replicative period observed when a microbial population is introduced to a new nutrient-rich environment. Its duration can have a pronounced effect on population fitness, and it is often measured in laboratory conditions. However, calculating the lag phase length may be challenging and method and parameters dependent. Moreover, the details of these methods and parameters used throughout experimental studies are often under-reported. Here we discuss the most frequently used methods in experimental and theoretical studies, and we point out some inconsistencies between them. Using experimental and simulated data we study the performance of these methods depending on the frequency of population size measurements, and parameters determining the growth curve shape, such as growth rate. It turns out that the sensitivity to each of these parameters depends on the lag calculation methods. For example, lag duration calculation by parameter fitting to a logistic model is very robust to low frequency of measurements, but it may be highly biased for growth curves with low growth rate. On the contrary, the method based on finding the point where growth acceleration is the highest, is robust to low growth rate, but highly sensitive to low frequency of measurements and the level of noise in the data. Based on our results, we propose a decision tree to choose a method most suited to one’s data. Finally, we developed a web tool where the lag duration can be calculated based on the user-specified growth curve data, and for various explicitly specified methods, parameters, and data pre-processing techniques.

**GRAPHICAL ABSTRACT:** 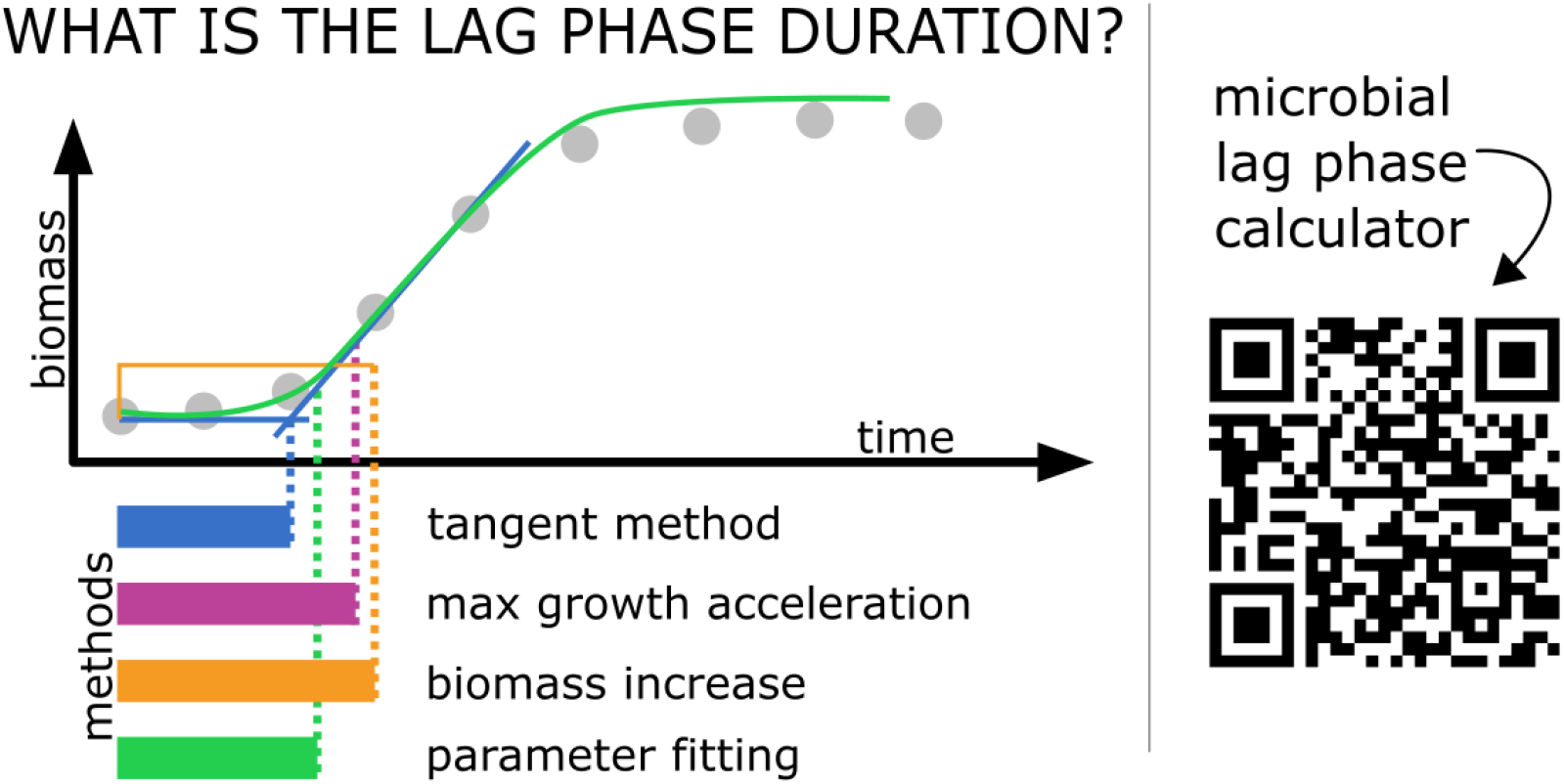

## INTRODUCTION

Microbial planktonic populations grown in a batch culture follow a predictable pattern in terms of how the population size changes in time. Such growth kinetics can be represented by a growth curve which is typically divided into the following phases: (1) lag phase, when cells adjust to a new environment before they start dividing, (2) exponentially growing phase (or logarithmical growth), when cells divide with a maximum rate and population’s density doubles regularly, (3) stationary phase, when cells cease divisions due to nutrients depletion, and, if the measurements are conducted long enough, the decline/death phase when population’s density drops due to the cell death. Such pattern of growth is typical for planktonic prokaryotic and eukaryotic microorganisms.

The lag phase was first described in 1895 by Müller as a temporary non-replicative period after bacteria are introduced to new media [1]. Indeed, adjustment to a new environment after growth-arrest requires broad cellular reorganizations, which are universal for prokaryotes and eukaryotes. The cells need to adjust their transcriptome, and proteome (e.g. in *S*.*cerevisiae* [2–4]) and rearrange cellular components that are necessary for nutrient uptake and biomass accumulation. These processes are activated shortly after environment change - for example, it was reported that gene expression in *Salmonella enterica* bacteria was affected as soon as 4 minutes after transfer into a fresh media, and within 20 minutes, almost a thousand genes were upregulated [5]. Proliferation restart requires induction of a broad range of processes, which include: glycolysis, nutrient sensing, amino acid metabolism, nucleotide biosynthesis, protein processing (including translation, folding, modification, translocation, and degradation), coenzyme and cell wall biosynthesis, transcription of genes involved in stress response, respiration, cell cycle control, and division. For more details about biological processes during the lag please see the review by Bertrand (2019) [6].

While many studies focus on the exponential phase and use the exponential growth rate as a measure of population fitness [7], the quantification of the lag duration is equally important to assess the stress or fitness of microbial populations [8,9]. Classically, fitness is defined by the number of progeny, which in microbiology is translated into the growth rate (i.e. the rate at which the population size doubles). However, the lag phase duration also affects fitness because it shows how quickly a given cell or population can adapt to an environmental change. Shorter lags enable earlier divisions, which might allow a cell to produce higher number of progeny within a fixed period of time. Such growth dynamics would be especially important in case of competition for limited resources or a limited time frame when resources can be used. Thus, the short lag phase is generally believed to be beneficial, and therefore populations in favourable conditions may be expected to evolve toward decreased lag duration [10]. However, the opposite strategy was observed in presence of antibiotics. In particular, when bacterial populations were exposed to antibiotics before a transfer to new growth media, they evolved towards increased lag duration which matched the duration of antibiotic exposure [11].

The variation in lag phase duration can be influenced by genetic, epigenetic, and environmental factors (e.g. [12–14]). For *Saccharomyces cerevisiae*, both nuclear and mitochondrial genes are important during the lag phase, and their expression patterns correlate with the lag phase length [13]. Besides genetics, it has been shown that the lag phase duration is history-dependent. Namely, previous cells’ exposure to given conditions shortens the time needed to adapt if the conditions are reintroduced [14–17]. What’s even more interesting - this effect has also been observed in daughter cells that had never experienced the initially introduced conditions what suggests epigenetic inheritance. [15]. Lags of individual cells may differ even within a clonal population grown in unaltered conditions. For example, older cells experience longer lags than their younger clones [18,19]. Such heterogeneity may be beneficial for the population, as it enables a bet-hedging strategy where a fraction of cells assures fitness advantage by fast growth in favourable conditions, while the other fraction provides survival in stressful conditions by extending their lag time [19–21].

There are multiple definitions of the lag phase and multiple ways of measuring its duration. One common definition is based on microscopic observations and it uses the first morphological signs of cell division as markers of the lag phase end (e.g. [15,22,23]). Within the studies on population-level, the lag phase can be defined as the time before any detectable increase in the cell abundance (biomass) [19], or as the time delay before a population reaches exponential growth [5]. To measure how the population’s density (cell abundance) changes in time standard laboratory methods can be used. Usually, one of the following methods is applied: spectrophotometry, colony counting on agar plates (CFU, Colony Forming Unit), and flow-cytometry. Viability counts (CFU) provide precise estimates of cell abundance even in a small population, however, this technique is time-consuming, requires previously determined culture dilutions, and is difficult to automatize and scale up. Nevertheless, due to its high sensitivity, it’s broadly used in food safety control [24,25]. Flow cytometry has a broad detection spectrum and it enables simultaneous measurements of various cell properties e.g. cell size or DNA content [26]. Spectrophotometry (optical density (OD) or absorbance) has narrower detection limits, but it is convenient, fast, cost-effective, and can be easily adapted for high-throughput testing via automatic measurements in constant time intervals.

Spectrophotometry is currently the standard way of obtaining microbial growth curves. However, one of its limitations is the fact that optical density is an indicator of biomass rather than cell counts (discussed by Swinnen *et al*. (2004) and Rolfe *et al*. (2012) [5,27]). To add to this problem, the optical density may be also affected by dead or lysed cells, or even by the cell shape which may change during the lag phase [28]. Additionally, if the inoculum size is small, exponential growth may start before the biomass reaches the OD detection level, and thus the observed (apparent) lag duration may be overestimated. This problem has been tackled by Baranyi (1999) [29] who proposed a so-called time to detection (TTD) method which has been applied in some experimental studies (e.g. [30]). The problem of initial biomass being below the detection level has also been tackled by Pierantoni *et al*. (2019) [31] who have proposed a new calculation method that uses the apparent lag phases to measure the initial population biomass. More broadly, the problem of density-based detection of growth for small populations could be mitigated by assuming certain model growth curve shapes below the OD detection level and/or by knowing the exact starting density of cells capable to proliferate (excluding dead and senescent cells) [32].

Mathematical models can be used to overcome some methodological limitations. In order to know when exactly cells started duplicating (i.e. the end of the lag phase), we would need to continuously monitor the number of cells. This is, however, not possible with methods such as spectrophotometry. As outlined above, the spectrophotometry measurements not only are taken in intervals, but also may be inaccurate if the cells change their mass or shape, or if their initial amount is below the detection level [5]. In such cases, some assumptions are needed to calculate the lag phase duration with sufficient accuracy. If one assumes that there is no population growth during the lag phase and then cells start synchronically dividing at a constant growth rate, the end of the lag phase can be calculated as the intersection between the tangent line to the point of maximum growth rate and the y = log(N0) line, where N0 is the inoculation density (hereinafter “tangent method” [6]; see for example: [12,15,24]). This is in fact the most frequently used method of calculating the lag duration. However, there are also other methods, for example: defining the end of lag as the point of the growth curve where the second derivative of the population size in time is maximal (hereinafter “max growth acceleration”, e.g. [33,34]), determining when the biomass increased from the initial value by some predefined threshold (minimal detectable increase, hereinafter “biomass increase”, e.g. [19]), or fitting experimental data to a mathematical model (hereinafter “parameter fitting to a model”, e.g. [8]). Various mathematical models have been proposed to account for the lag phase [27], and there are tools and packages which use those to estimate the lag duration from the experimental data (for example R package *nlsMicrobio* [35]). However, none of these tools is strictly focused on calculating the population lag duration. Moreover, they require a good knowledge of the models and R programming skills, which may make them difficult to use. Finally, as discussed in Baty *et al*. (2004) [32], the data quality impacts the lag duration measurements to a higher extent than the choice of a model. Although Baty *et al*. (2004) [32] investigated the insufficient number of data points as a potential problem in lag duration calculation, experimental biologists may face other problems with the data quality such as noisiness or growth curve shape that deviates from mathematical models [36]. Interestingly, technicalities related to dealing with such ‘unideal’ data tend to be omitted in methodologies described by empirical studies that measure lags. They have also not been discussed in theoretical studies which focus on the mathematical formulations and biological assumptions rather than the reality and limitations of laboratory experiments. These facts make a knowledge transfer between theoretical and experimental biologists difficult.

Within this study, we describe the methods most frequently used to measure the lag duration and discuss how these methods stem from various mathematical models describing microbial growth curves. We highlight the advantages and limitations of each lag calculating method. We further use both empirical and simulated growth curve data to show how the lag duration estimate may depend on the data quality (i.e. noisiness), the shape of the growth curve, and the lag duration calculation method. We propose a decision tree that may be useful in choosing the lag calculating method best suited to one’s data. We aim to emphasise that chosen methodology, parameters, and data pre-processing can strongly influence the results, which is especially visible for less typically shaped growth curves. Finally, we develop a publicly available web server MICROBIAL LAG PHASE DURATION CALCULATOR (https://microbialgrowth.shinyapps.io/lag_calulator/) which allows calculating the population lag duration according to various state-of-art methods. The calculator allows for fast and easy data analysis and direct comparison of different methods.

### The lag phase in mathematical models

There are many mathematical models that describe the microbial population growth depicted by a growth curve. They include the traditional simple models such as exponential [37], logistic [38] or Monod growth [39], as well as more modern stochastic or agent-based models (see the review by Charlebois and Balázsi (2019) [40]). While the pure exponential model is parametrised only by the *growth rate*, the other models may depend on multiple parameters. For example, the logistic model introduces the *carrying capacity* parameter which represents the maximum population size, and Monod model introduces the so-called *half-velocity constant* (which quantifies the relationship between nutrient concentration in media and population growth rate). Another version of this model may be parametrised by the *maximum resource uptake rate* and the *saturation constant* as derived by Michaelis-Menten.

Please note, that within this publication "method” refers to the way in which the lag phase is determined, while “model” refers to a set of equations that reproduce the entire microbial growth curve.

Out of the three typically described growth phases (lag, exponential and stationary phase), the exponential and stationary phases are recovered by most of the models discussed in Charlebois *et al*. (2019) [40]. One exception is the pure exponential model which only describes the exponential phase. However, the lag and death phases require additional assumptions and are rarely described with sufficient detail. The models that aim to capture or predict the lag phase are well summarised in the two extensive reviews [27,32]. Some models describe the lag only phenomenologically. For example, the Baranyi (1993) and Baranyi and Roberts (1994) [41,42] models assume that there is some adjustment function that describes the population’s adaptation to a new condition. On the other hand, there are models that attempt to make specific assumptions regarding what happens in the lag phase. The Hills and Wright (1994) [43] assume that certain biomass needs to be reached for the cell to start the chromosomal replication. Consequently, their model independently tracks in time the amount of biomass and chromosomal material. The model proposed by McKellar (1997) [44] assumes there is some heterogeneity in the population, and that one part of the population grows exponentially from the very beginning, whereas the other part does not replicate. A more realistic version of this model was proposed by Baranyi (1998) [45] where the non-replicating cells are assumed to transform at a constant rate into the replicating ones, meaning that they exit the lag phase. Finally, a model by Yates (2007) [46] adds another compartment, namely cells that die at a constant rate. This specific assumption allowed the model to reproduce the initial biomass decline which is sometimes observed during the lag phase. Interestingly, as noticed by Baty (2004) [32], some of the models described above are equivalent, in spite of being based on distinct biological assumptions.

### The lag phase calculation methods

The most popular and intuitive method of calculating the lag duration (“tangent method”) defines the lag phase end at the intersection point of log(N0) (where N0 denotes the initial biomass) and a line tangent to the logarithm of population size in the exponential phase [6,39]. Why bother about more complicated mathematical models?

The definition above is in fact a consequence of the pure exponential growth assumption. If cells do not divide for some time λ and then start growing with a constant growth rate, then the time λ perfectly corresponds to the lag as measured by the intersection between the lines described above (Table 1, Fig. 1: exponential model & tangent method).

| Name | Lag end calculation method | Underlying model assumptions | Challenges | Example data for which the method is suitable | Examples of experimental studies using the method |
| --- | --- | --- | --- | --- | --- |
| BIOMASS INCREASE | The increase of OD or biomass by a certain value from the beginning of the growth. | The population size does not increase its biomass during the lag and then starts growing with any growth rate that may be variable or hard to measure. The increase by the threshold value is the minimal increase possible to detect with high confidence. | Choice of the threshold value is arbitrary (not 'biologically- relevant') | Data where the initial size of the viable population cannot be determined with high confidence. | Opalek *et al.* 2022 [19] |
| MAX GROWTH ACCELERATION | The point of the growth curve where the second derivative of population size in time is maximal. | The population size does not increase its biomass during the lag and then starts growing with a decreasing growth rate. | Noisy data may affect the detection of such a point. | Growth in a batch culture where the amount of resources decreases with time and the amount of unfavourable by-products increases with time. | Liu *et al.* 2021 [33] |
| TANGENT METHOD | The intersection of the initial density line and the line tangent to the growth curve at the maximal growth rate (on log. scale). | The population size does not increase its biomass during the lag and then starts growing exponentially with a constant growth rate. | If the growth rate varies, it may be challenging to find the "real" or maximal growth rate. Moreover, the initial density needs to be determined with high confidence | Growth in constant resources and environmental conditions (e.g. chemostat (at starting point), serial transfers). | Valík *et al.* 2021 [24] |
| PARAMETER FITTING TO THE BARANYI MODEL | Using parameter fitting procedures in order to simultaneously find all Baranyi & Roberts model parameters that provide the best fit to the entire growth curve. Then the lag is defined as $\ln(1 + 1/q_0)/r$ where $r$ is the maximal growth rate and $q_0$ represents the physiological state of the inoculum. | The population size grows according to the logistic model adjusted by function: $\alpha = q(t)/(1 + q(t))$ where $q(t) = q_0 e^{vt}$ describes the concentration of some critical substance. The adjustment function slows down the initial growth but it does not pause it during the lag time as assumed by other methods. | The lag duration fitted to the data may depend on many technical parameters of the fitting algorithms, or it may not be found if the fitting algorithms fail to converge. Finally, the lag understood by Baranyi is not the time when cells do not divide, but the time required to adjust to the new media (see Discussion for more details) | Data that is well described by the Baranyi & Roberts model, where ideally some parameters ($r$, $K$, or $q_0$) are known *a priori*. | Kim *et al.* 2022 [50] |
| PARAMETER FITTING TO THE LOGISTIC MODEL | Using parameter fitting procedures in order to simultaneously estimate the lag duration, carrying capacity, and growth rate parameter values. | The population size does not increase during the lag and then starts growing according to the logistic model: i.e. with the growth rate decreasing according to the following equation: $r(1-N(t)/K)$ where $r$ is the maximal growth rate, $K$ is the maximal population size, and $N(t)$ is the population size at time $t$. | Assumes a very specific growth curve shape. Moreover, the lag duration fitted to the data may depend on many technical parameters of the fitting algorithms. Finally, the fitting methods may not converge to any solution. | Data that is well described by the logistic model, where ideally some parameters ($r$ or $K$) are known *a priori*. | Reding-Roman *et al.* 2017 [8] |

**Fig. 1.**
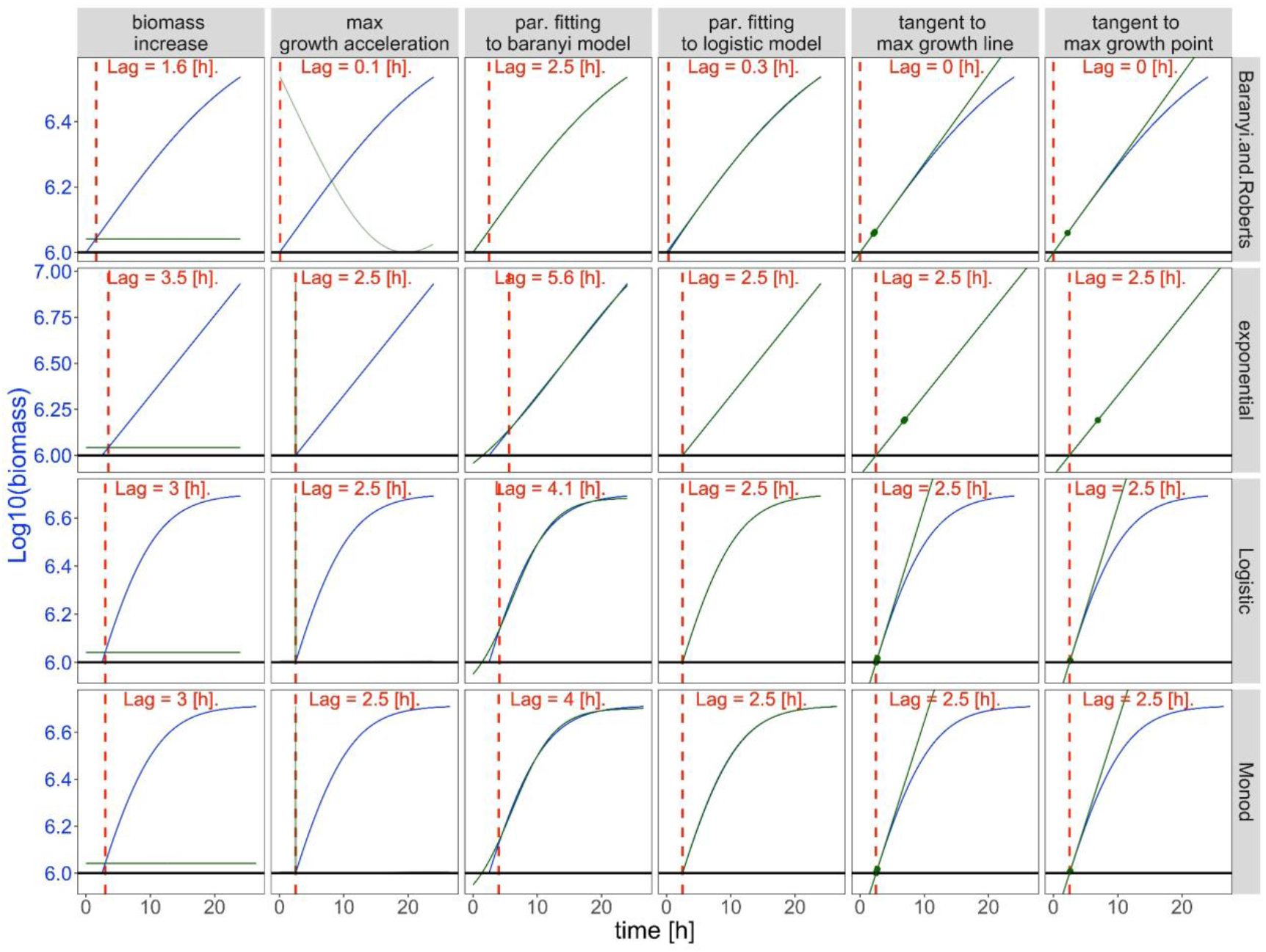
Lag phase durations (given in red) calculated for data (blue lines) simulated under various models (rows) and calculated by different methods (columns). The red dashed lines indicate the end of the lag phase, black solid lines indicate the initial biomass (first measurement) and the green lines are (left to right): the threshold of log(biomass) value at which the lag phase is assumed to end [biomass increase method]; second derivative of log(biomass) scaled to the values visualised in the plot [max. growth acceleration method]; fitted growth curve [par. fitting to the Baranyi model and par. fitting to logistic model methods]; tangent lines to the exponential growth [tangent to max. growth line and point methods].

However, microbial populations in a batch culture do not grow with a constant growth rate. According to the Monod model assumptions [39], the growth rate depends on the availability of the resources, and therefore it decreases over time. The growth rate may also depend on other factors, e.g. the population density [47]. In the case of cancer cells or in populations that reproduce sexually the growth rate may be lower at low population density, which is known as the Allee effect (e.g. [48,49]).

This is why, in order to assess the most exact time when cells start divisions, there is a need for a mathematical model that is likely to represent the empirical growth curve and to fit the lag time together with other growth parameters. In particular, note that when there is some growth detection threshold, the time by which we notice any growth (i.e. apparent lag) may be impacted not only by the lag length but also by other growth parameters. For example, when the initial biomass and growth rates are very low, the slow growth may not be detected and treated as lag. This problem has been extensively discussed in Pierantoni *et al* (2019) [31].

The summary of the most popular methods of calculating the lag duration, the assumptions underlying each of the methods as well as possible challenges related to each method are given in Table 1.

## RESULTS

### Testing methods for lag duration determination on the simulated dataset

Given that the methods of calculating the lag phase duration described above (Table 1) were based on theoretical assumptions, we first simulated microbial growth curves based on various well-known deterministic mathematical models: (i) the simple EXPONENTIAL model which assumes that microbes do not grow nor divide for some time (lag) and then start growing exponentially with a constant growth rate; (ii) LOGISTIC model which assumes the microbes do not grow nor divide for some time (lag) and then start growing exponentially with a decreasing growth rate; (iii) Monod model where microbes do not grow nor divide for some time (lag) and then the speed of the growth after the lag phase is coupled with the decrease in resources, and (iv) Baranyi model which assumes cells do grow and divide in the lag phase but that growth is slower than in the exponential phase. See the Appendix for the formulation of each model. We set that for all generated growth curves the lag phase lasts for 2.5 hours.

Then, for each of these simulated growth curves, we calculated the lag phase duration using the well-established methods described in Table 1. The most common tangent method was further split into: tangent to point (i.e. where the tangent line is drawn to the first point where the growth rate is maximal) [23] and tangent to line method (i.e. where the tangent line is understood as a regression line fitted to a number of points around those with the maximal growth rate) [7].

Max growth acceleration, parameter fitting to the logistic model, and both tangent methods found the correct lags (lag = 2.5 h) for data simulated under exponential, logistic, and Monod models (Fig. 1). Even the simplest tangent method worked well for the data simulated under these models, even though the data does not necessarily meet the assumption of a constant growth rate. Importantly the Baranyi model seems to be inconsistent will all other models. Namely, the lag durations calculated for the data simulated under the Baranyi model are underestimated by all methods apart from the one where parameters are fitted explicitly to the Baranyi model. Conversely, the parameter fitting to the Baranyi model method tends to overestimate the lags for data simulated under all models apart from Baranyi. The biomass increase method overestimates lag phase duration by one timepoint (0.5 h) for logistic and Monod models and two timepoints (1 h) for the exponential model. It is the consequence of this method formulation, where lag phase end is defined as the first timepoint after detectable growth.

### Testing methods for lag duration determination on the empirical dataset

Having verified how the lag calculation methods work on simulated data (Fig. 1), we set off to verify how these methods perform on real experimental data. We used 11 growth curves from *Saccharomyces cerevisiae* grown in various conditions (see Methods and Supplement). Some of these curves resemble the model data, while others are much noisier and challenge some of the model assumptions, for example: the biomass drops at the beginning of measurements (Fig. 2: curve_5 and curve_6), there is no typical exponential phase (Fig. 2: curve_11) or the lag phase is boldly prolonged and turbulent (Fig. 2: curve_7 and curve_11). Results obtained for empirical data show that the lag duration estimates may vary depending on the lag calculation method (Fig. 2, Supplementary Table 2). The lowest discrepancy between different methods is observed for curve_1, where the lag duration calculated by all methods is 1.86 ± 0.83 h. In some cases, all but one method give similar results (e.g. curve_7, where the biomass increase method gives an outlying result), in others, some methods fail to find the correct lag duration (e.g. curve_7, curve_11) due to the data noisiness (Fig. 2). The highest discrepancy between methods was obtained for curve_11 where a minimal lag phase length of 1.5 h was obtained by the max growth acceleration method, while both tangent methods and parameter fitting to logistic model yielded the lag duration of ∼18h.

**Fig. 2.**
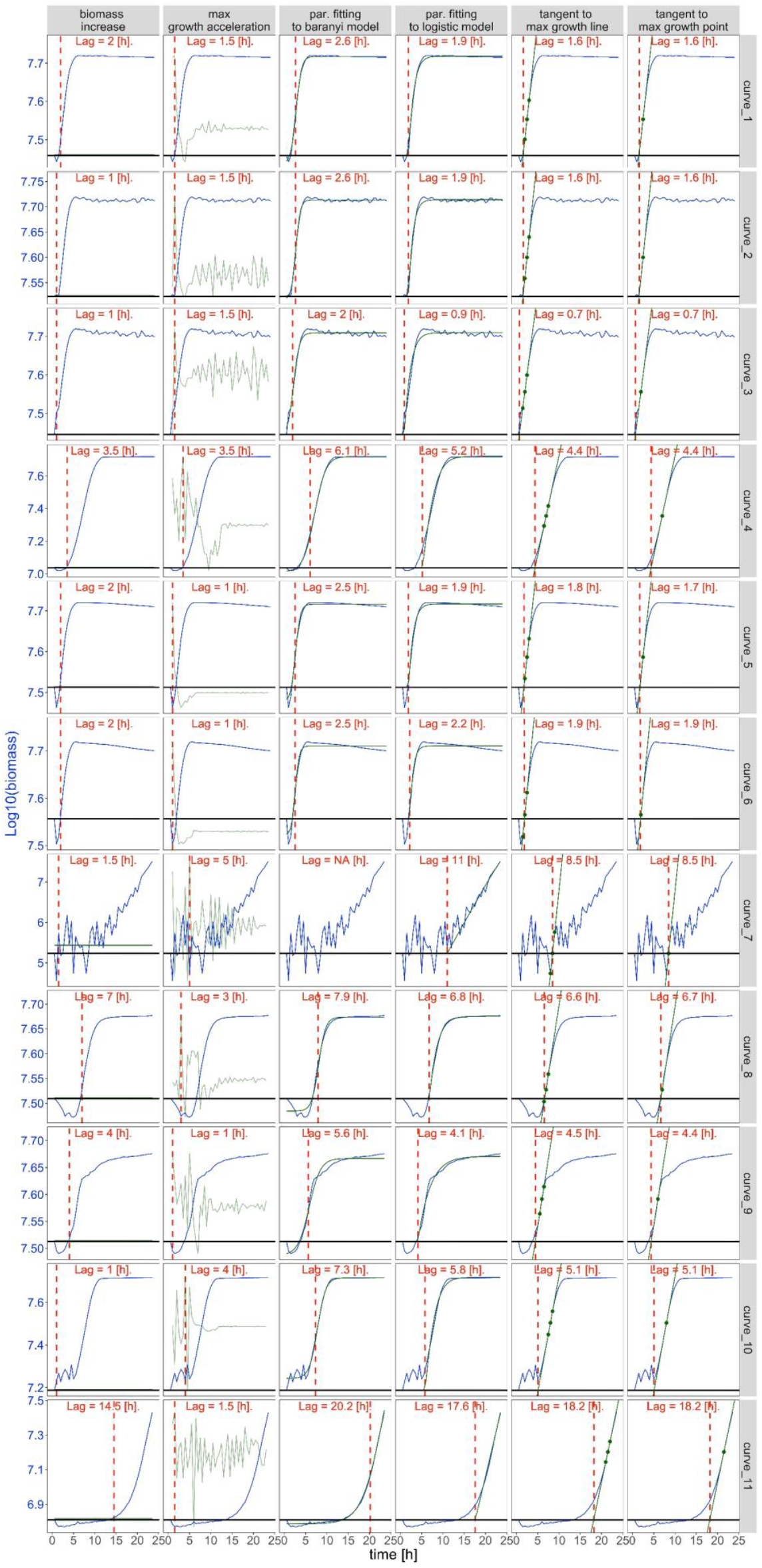
Lag phase duration (given in red) calculated for empirical growth curves (blue lines) obtained for *S. cerevisiae* grown in various conditions (top to bottom). The red dashed lines indicate the end of the lag phase, black solid lines indicate the initial biomass (first measurement) and the green lines are (left to right): threshold value at which the lag phase is assumed to end [biomass increase method]; second derivative of log(biomass) scaled to the values visualised in the plot [max. growth acceleration method]; fitted growth curve [par. fitting to Baranyi model and par. fitting to logistic model methods]; tangent lines to the exponential growth [tangent to max. growth line and point methods].

### Testing the sensitivity of lag determination methods to data noisiness

In order to understand the source of possible errors and biases in lag calculation on experimental data, we simulated growth curves from the logistic model (which in our experience the most adequately represents the real growth curves) and to each curve we added noise simulated from the random distribution with mean = 0 and standard deviation dependent on the initial biomass B0 (Supplementary Fig. 2). Thus, our simulated growth curves could be described as:

*B*_*noisy*_(*t*) = *B*(*t*) + *N*(0, *sd* * *B*(0)), where B(t) is the solution from the deterministic logistic model with a lag component (see Supplement for the formulation of this model).

We varied the level of noisiness (i.e. set *sd* between 0 and 0.5) together with other parameters such as: population growth rate, lag time, and time interval between data points (which represents the frequency of population size measurements). For each combination, we simulated 100 curves. Then for each of these curves, we calculated the lag according to each of the lag calculation methods and we calculated the bias i.e. the difference between observed and expected lag.

First, we tested how the frequency of population size measurement impacts the lag estimation. In agreement with previous reports [32], it turned out that frequent measurements improve the accuracy of lag duration estimation. This effect can be observed within all lag calculation methods (Fig. 3: top vs bottom row) and it is especially pronounced for data with high noise (sd = 0.5). Large time intervals between data points (Fig. 3: bottom row) lead to increased variance in the lag duration estimation (wide boxplot) and to high bias (median value deviates from y = 0 line). It is especially pronounced in the biomass increase and max growth acceleration methods which overestimate lag durations even when there is minimal amount of noise (low sd). Such bias results from the fact that these methods operate only on the data points provided and they do not use any implicit models to interpolate between them.

**Fig. 3.**
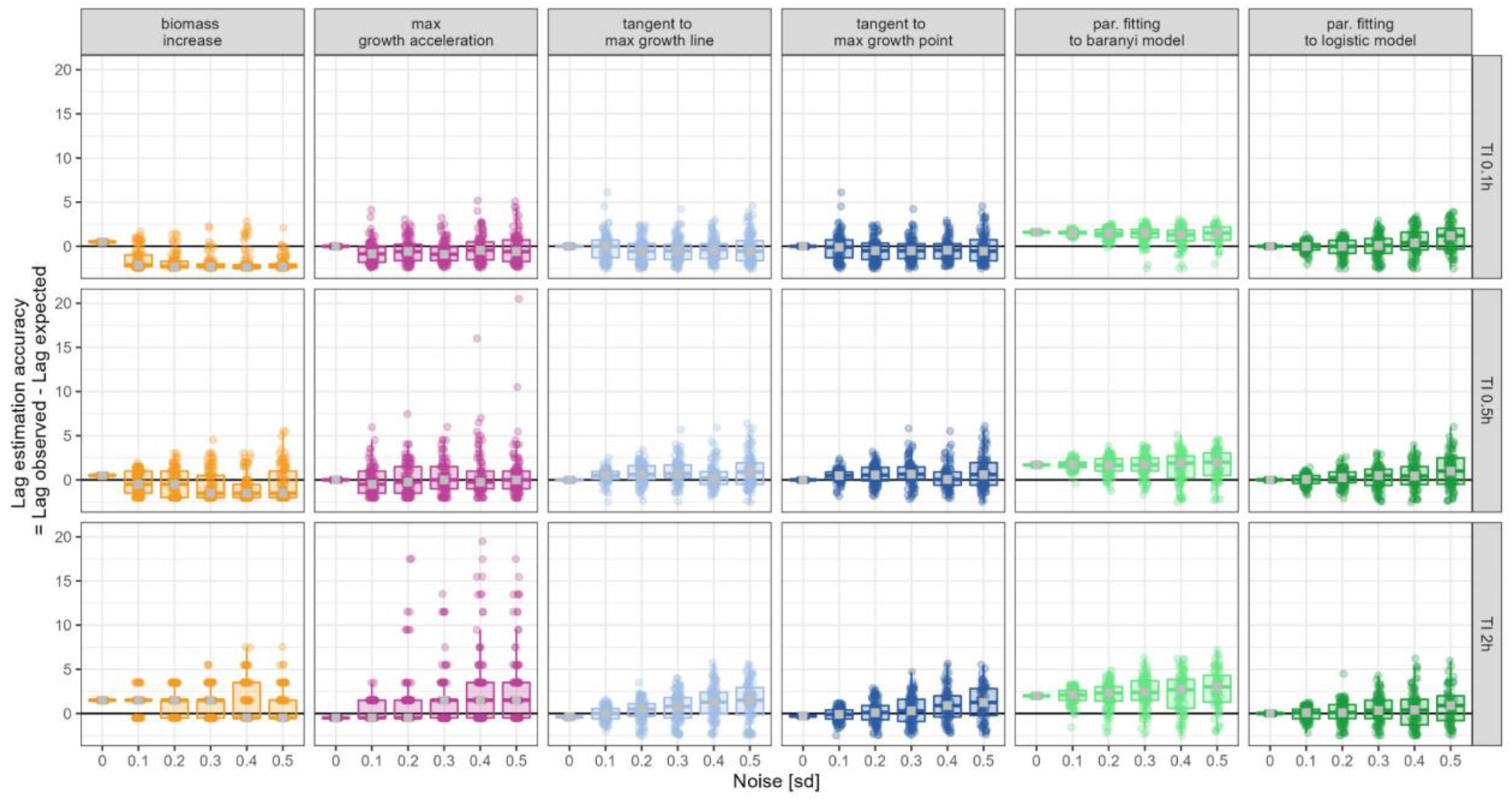
The impact of time intervals (TI) between subsequent measurements of population size on lag phase duration estimations. For each sd (noisiness, x-axis), measurement frequency (time interval, rows), and lag calculation method (columns) 100 simulations were conducted (points). The y-axis illustrates the difference between true (expected, set as 2.5 h) and estimated (observed) lag. Boxplots illustrate the distribution of points, and the median value (grey square). All points located above y = 0, show these calculations where lag phase length was overestimated, similarly, all points below y = 0 show calculations where lag duration was underestimated.

Next, we checked whether the population’s growth rate has an impact on lag estimation. We calculated lags for three growth rates representing slow, moderate, and fast growth, and for varied levels of data noisiness (Fig. 4). For most of the methods slow growth results in low accuracy of lag estimation, which may additionally decreases when data is noisy. These results are consistent with our experience, namely, when the growth curve is flat, it is challenging to find a point where the population starts growing exponentially (Fig. 4: top row). In contrast, even for noisy data, fast growth facilitates lag phase estimations (Fig. 4: bottom row). The only method which is not affected by the growth rate is the biomass increase method, however it systematically underestimate lag duration, and this bias increases with introduced noisiness. Conversely, the parameter fitting to the Baranyi model systematically overestimates lag lengths especially when the population grows slowly. Max growth acceleration and both tangent methods are not biased, and are sensitive to growth rates to a similar degree, where lag estimates becoming less accurate with increasing noise. Interestingly, parameter fitting to the logistic model shows good accuracy and no bias for moderate and fast growth, but it suffers from low accuracy and high bias in the case of low population growth rate and high levels of noise.

**Fig. 4.**
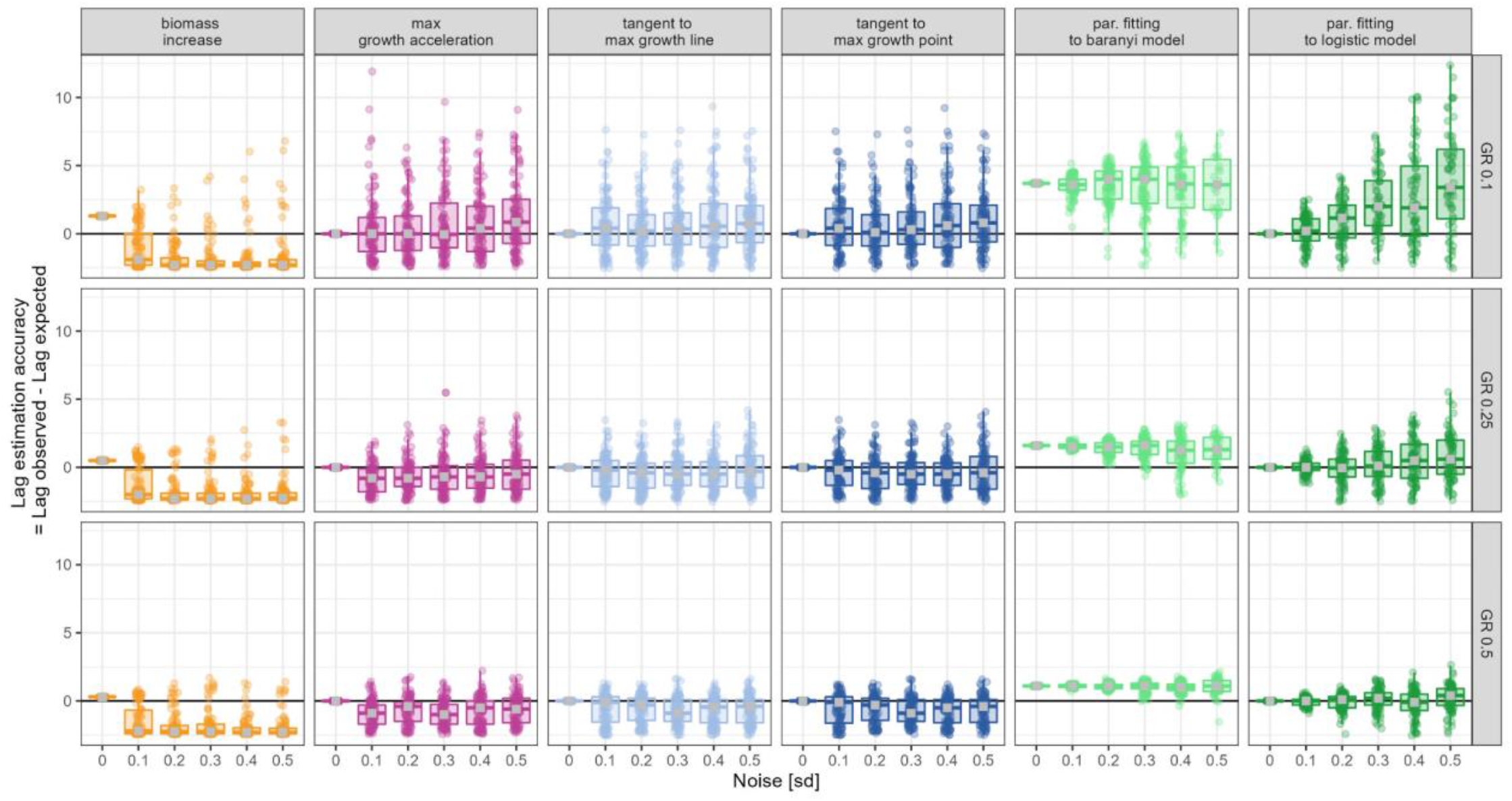
Impact of growth rate (GR) on lag phase duration estimations. For each sd (noisiness, x-axis), growth rate (rows), and lag calculation method (columns) 100 simulations were conducted (points). The y-axis illustrates the difference between true (expected, set as 2.5 h) and estimated (observed) lag. Boxplots illustrate the distribution of points, and the median value (grey square). All points located above y = 0, show these calculations where lag phase length was overestimated, similarly, all points below y = 0 show calculations where lag duration was underestimated.

Another factor that affects the lag length estimation accuracy is the actual length of the lag phase. The longer the lag, the easier it is to erroneously detect some noise during the lag phase as the first signs of growth. Indeed, if a population has a long lag phase, its duration tends to be underestimated, while very short lags tend to be overestimated by all methods. This bias is increasing with higher noisiness (Supplementary Fig. 3), with the estimations of lag phase duration by parameter fitting to the logistic model being the most robust.

The comparison of biases of all lag calculation methods are shown in Fig. 5. The methods that tend to be systematically biased within our parameter range are the biomass increase (underestimates the lags) and fitting to Baranyi model (overestimates the lags). The biomass increase tends to erroneously pick random points as sudden lag phase end, which leads to underestimation of lag phase duration. Conversely, the parameter fitting to the Baranyi model systematically overestimates lag duration. This is a consequence of the dataset being simulated by the logistic model which is inconsistent with Baranyi’s assumptions. The other methods are less biased within our parameter range. Both tangent and max growth acceleration methods have poor performance on growth curves with long lags, and max growth acceleration method is very sensitive to high levels of noise. The parameter fitting to the logistic model is the most robust to data noisiness, however, interestingly what affects it the most is the low growth rate.

**Fig. 5.**
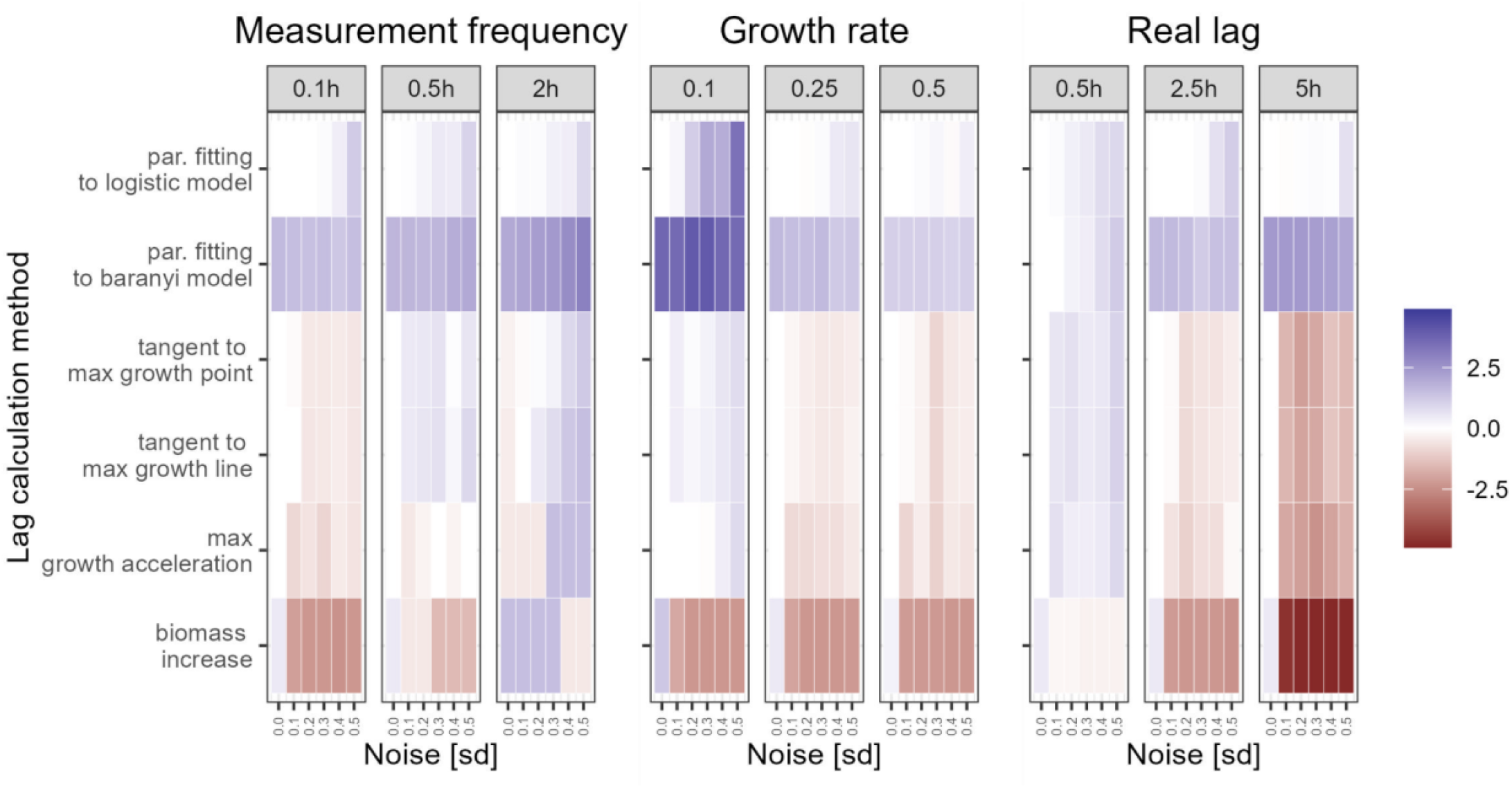
The median bias of lag estimation of each of the lag calculation methods applied to growth curves with varied level of noise (x-axis) and other growth curve parameters (column panels). White squares mean the method is unbiased, blue means the lags tend to be overestimated, and red means the lags tend to be underestimated.

### Testing the effects of data pre-processing on estimated lag lengths

Lag estimation is challenging if data is noisy. This problem could be mitigated by some data pre-processing techniques that allow for removing noise from the data. Therefore, we verified how manual curation or data pre-processing affect the lag calculation accuracy. We applied the following pre-processing methods to our experimental curves (Fig.2): (i) smoothening the curve by Tukey’s smoothing function, (ii) cutting the data at 12h to remove the noise observed in the stationary phase. It turned out that for the “typical” growth curve shapes, such as the ones presented in curve_1 and curve_2 (Fig. 2), various algorithms are consistent and the data pre-processing does not influence the results. However, the noisier the data, the less consistent the results based on multiple lag calculation methods and data pre-processing techniques (Fig. 3). We have additionally verified that adding a constant to the entire growth curve (e.g. by neglecting the blank correction) affects the lag phase length estimates given by all but the biomass increase method (Fig 3). Importantly, such a constant value added to the growth curve may not only relate to blank correction, but also to the case when a fraction of the population is dead or damaged and does not duplicate throughout the growth. We further investigate how that phenomenon affects the lag calculation in Supplementary Fig 1.

### How to choose the lag calculation method best suited to one’s dataset?

We propose a decision tree to facilitate the choice of lag calculation method (Fig. 7A). The recommendations are based on our results shown in previous sections. Altogether, we suggest trying to estimate lag duration by parameter fitting to the logistic model in the first place. This method is the most robust, it captures whole growth dynamics and because of that, it mitigates technical limitations (such as a device’s detection limits). On the other hand, the biomass increase method is the least dependent on any assumptions. In particular, it is the only one that is not affected by the blank correction or existence of dead cells in the culture. Therefore we recommend it if the other methods cannot be applied. Additionally, we encourage to use multiple methods and to investigate possible inconsistencies between their results.

**Fig. 6.**
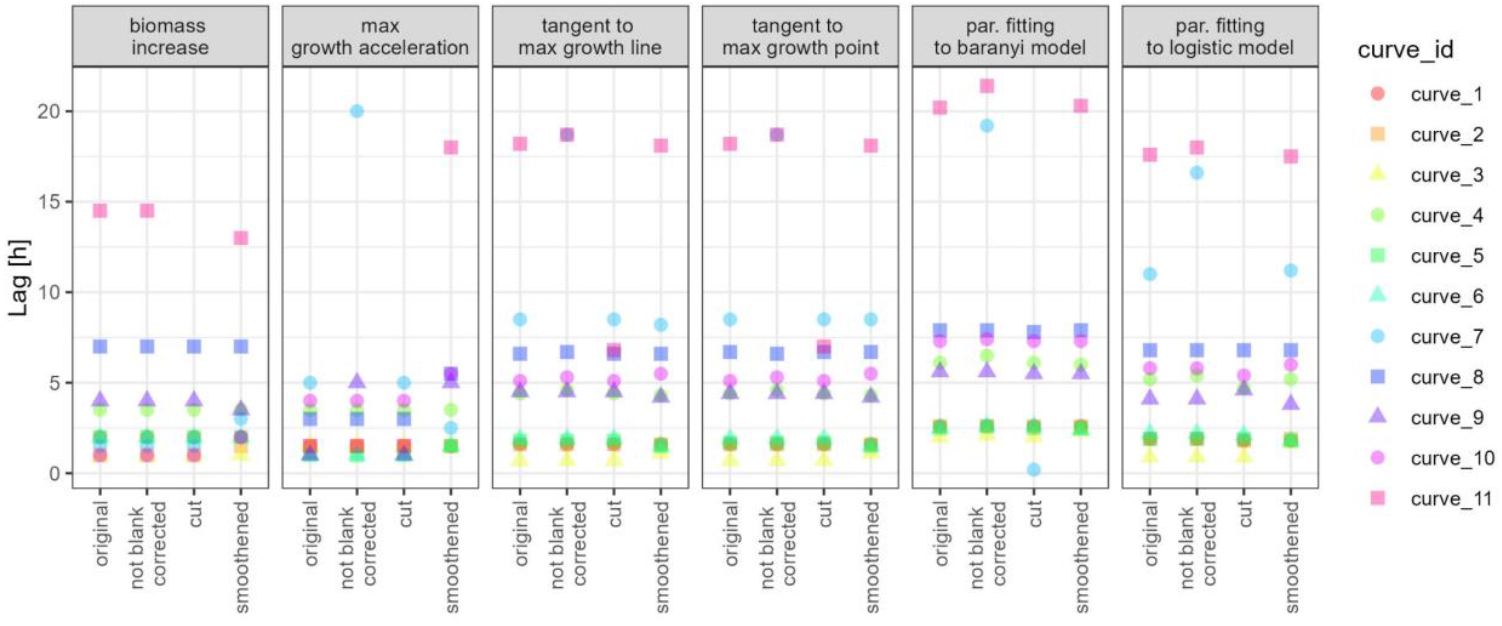
Distribution of lag duration estimated by six discussed methods (columns) per each curve (colour & shape) and pre-processing algorithm (x-axis). “Original” means no pre-processing has been applied, “not blank corrected” means data were not corrected for the blank value, “cut” means the data curve has been shortened to 12 hours only, and “smoothened” means Tukey smoothening has been applied.

**Fig. 7.**
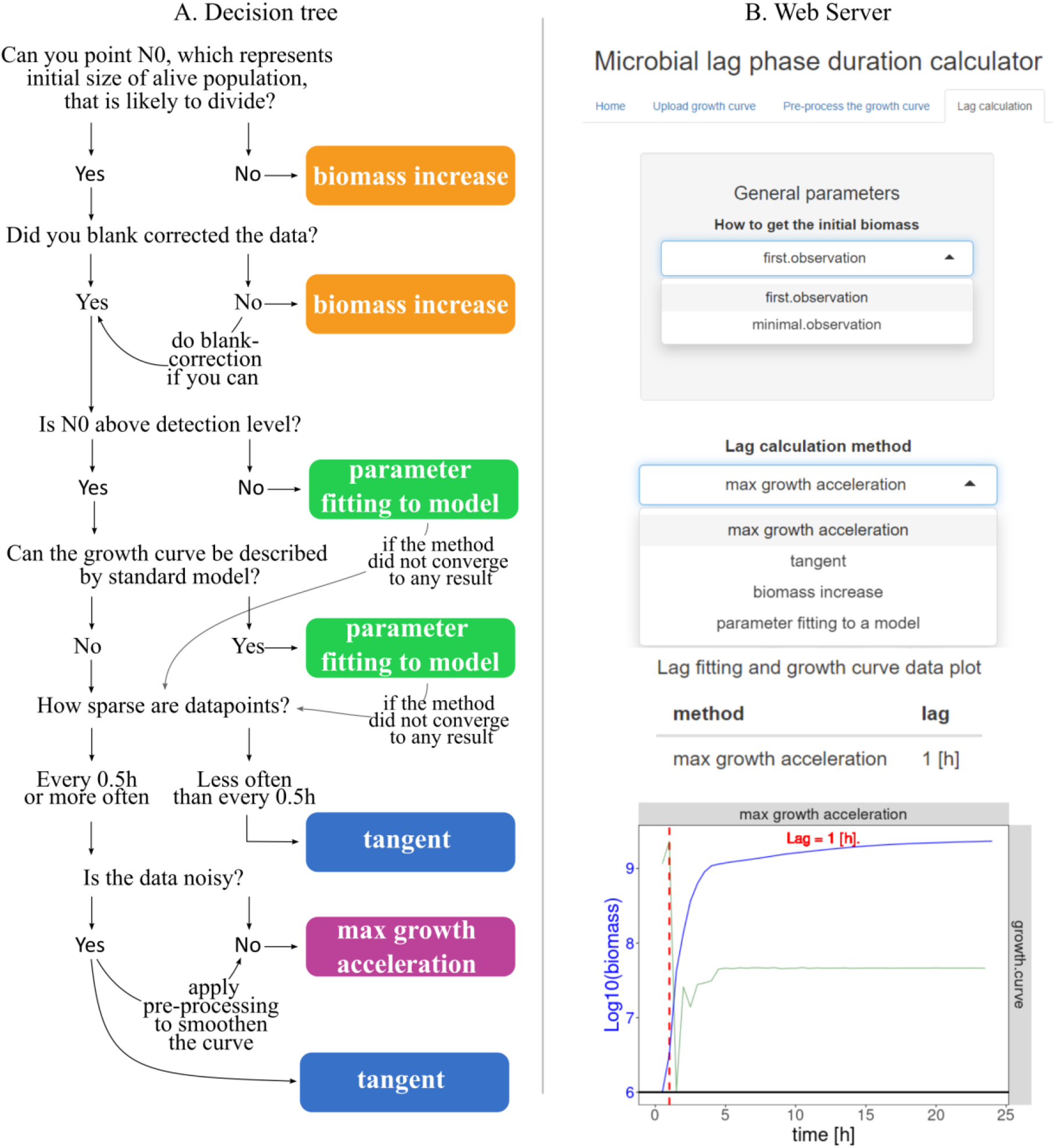
A: Decision tree to facilitate the choice of appropriate lag calculation method. B: Print-screen from web server MICROBIAL LAG PHASE DURATION CALCULATOR where lag phase duration can be calculated for a user-specified growth curve, and by any of the methods discussed in Table 1.

### The lag phase duration calculator

Finally, in order to allow other researchers to calculate lag duration by different methods and compare the results, we created a web server MICROBIAL LAG PHASE DURATION CALCULATOR, where lags can be calculated by various methods upon insertion of the growth curve data (i.e. table with time and biomass columns). The webserver is freely available under the following address: https://microbialgrowth.shinyapps.io/lag_calulator/

The MICROBIAL LAG PHASE DURATION CALCULATOR is designed to automate the lag duration calculation process. The tool does not require any programming, nor mathematical modelling skills. Additionally, we deposited our code and lag calculating functions on GitHub so that they can be used locally, customised, and further improved.

## DISCUSSION

Lag phase duration is an important fitness component for microbial populations. Within this publication, we discussed the most popular approaches to calculate the lag phase duration for population-level data. We implemented these approaches and tested them on simulated and empirical data to show that they may disagree depending on the shape of the growth curve. To further compare the methods in terms of their accuracies and biases, we simulated growth curve data with varied level of noise and other parameters such as growth rates, lag lengths or the time intervals between data points (understood as the frequency of measurements). For each of these growth curves we estimated the lag duration with each of the lag calculation methods. It turned out that the noise and other parameters affect the quality of lag duration estimation, hence we tested how smoothening the growth curve and other data pre-processing techniques influence the lag estimation. Since various methods were sensitive to various parameters (for example the max. growth acceleration method was largely affected by the high noise, while the parameter fitting to logistic model method was affected by low growth rates), we used our results to propose a decision tree designed to help in choosing the lag calculation method best suited to one’s data. We have also developed a web tool where the lag duration can be calculated based on the user-specified growth curve data, and for various explicitly specified methods, parameters, and data pre-processing techniques. Additionally, we deposited our code on GitHub so that the function “Calculate.Lag” can be taken directly to the R environment for local use and customization.

Here, we chose the four most frequently used approaches to estimate lag phase duration. We compare: THE BIOMASS INCREASE, which uses a predefined value of a biomass (or absorbance) gain that can be confidently marked as population’s growth; (ii) THE MAX GROWTH ACCELERATION, which assumes the lag ends at the time point where the second derivative of a population size in time is maximal, (iii) THE TANGENT METHOD, where lag phase endpoint is marked as the intersection point of a line tangent to maximal growth and a line y = log(N0), where N0 is population density at the inoculation; and (iv) PARAMETER FITTING TO A MODEL which simultaneously finds best-fitted parameter values for the entire curve (e.g. lag phase length, maximal population size, maximal growth rate) using logistic or Baranyi models. All these methods are developed based on typically shaped growth curves. Therefore, we tested the accuracy of their estimations using simulated (typical) and empirical (typical and untypical) growth curve data. As expected, for data with typically shaped growth curves, the majority of the methods showed similar results. This is in line with the previous observation, that the choice of a model influences the calculated lag phase duration to a lesser extent than the data quality and characteristic [32]. Interestingly the Baranyi model [51] is not consistent with other methods. This is a consequence of how the lag is defined within Baranyi model. While other models assume the lag is the time when the population size does not grow, Baranyi defines lag as the delay between the population size expected if the cells started growing immediately after inoculation with the maximum growth rate and the observed one [32]. This is consistent with the original definition [52] which defines lag duration as the difference between the time expected to take to reach the observed population size if population size was growing at a maximal rate and the actual time taken to reach that population size (including the lag duration). Note, however, that such definition is affected by all factors that lead to a slower growth (e.g. nutrient depletion), and may be dependent on the time when measurements are taken.

Taken together, the lag phase estimation cause little or no trouble when the growth curve resembles a standard model shape, and in such cases, the choice of the lag calculation method can be driven by one’s preferences. It, however, becomes more complicated for noisy or untypical growth curves.

In line with previous research [32], we demonstrated that the frequency of measurements can strongly influence the lag phase duration estimates (Fig. 4). We recommend taking measurements with maximal 0.5h intervals, and more frequently if one expects untypically shaped growth curves. We also highlight the importance of correct calculation of N0 (the initial number of alive cells, capable of proliferating) and of that number being above the detection limit. In the laboratory settings, this number can be relatively easily adjusted by adequate dilutions and simultaneously gives a much broader spectrum of possible analysis afterward. If N0 is below the detection level, then we are likely to overestimate the lag duration, because the first signals of growth will also be under detection level. To overcome this problem, one can assume a certain growth curve shape below the detection limit as done in Pierantoni *et al*. 2019 [31] and apply model fitting to estimate the lag duration. Additionally, if this number cannot be measured with high confidence (for example because of some dead or senescent cells being a part of the inoculum) one can use the biomass increase method to estimate population lag duration.

Although THE BIOMASS INCREASE method is simplistic, and it may be questionable if its results represent the real lag length, we believe it provides a good ecological measure of how efficiently a given population can inhabit a niche (net biomass gain). We suggest to apply this method if the growth curve greatly deviates from model shape or when the growth curve cannot be corrected for blanks or dead cells (i.e. if a fraction of the population size accounts for dead cells). In this case all other methods do not work correctly, because their assumptions are violated (Fig. 6). Note however that an important drawback of this method is that the chosen threshold value is arbitrary and may have no biological meaning.

The max growth acceleration is a very elegant method from mathematical point of view, and it may be a good choice for growth curves with non-standard shapes. However, it is very sensitive to noise (i.e. the calculation of the second derivative is very noise sensitive, Fig. 5). We recommend using smoothening function before applying the max growth acceleration method (Fig. 6).

The most popular TANGENT METHOD works reasonably well even if the assumption about the constant growth rate is violated. Additionally, it is not highly affected by data noisiness, but it performs worse than the parameter fitting method if the measurements are not taken often enough (Fig. 5). The challenging step may be the choice of how to draw the tangent line. If the tangent line is drawn to one point only (the point where the growth rate is maximal) there is a risk its slope will be under or overestimated if the data is noisy and an outlying point is chosen. This problem can be mitigated by drawing the regression line around points in the exponential phase. However, it may be not obvious in which time range the population grows exponentially. In fact, in order to know where the exponential phase starts one needs to know where the lag phase finishes, which brings us back to the original problem. Thus, the selection of data points in exponential phase often requires some manual inspection or additional assumptions. Within our web tool, N points are chosen around the point with the maximal growth rate, where N is a user-specified parameter. Note that the tangent method requires the initial number of cells capable of proliferating (N0) to be determined with high confidence. If a substantial fraction of the population is dead or senescent, what may be the case for populations that had previously experienced some stress (e.g. drug exposure, long-term starvation) [19,36], the N0 captured by absorbance will be heavily overestimated.

The parameter fitting to the logistic model is the most robust lag calculation method. It shows good accuracy even for noisy data (Fig. 5), and it can be used to overcome some technical limitations (such as N0 below detection level, or a high level of noise). Importantly, the method performed well not only for the curves simulated from logistic model, but also for the ones simulated from Monod model (Supplementary Fig. 4). The performance of this method depend on the shape of the growth curve, e.g. if the growth curve highly deviates from the standard shape, the fitting may not converge to any solution (Supplementary Fig. 5). This problem can be fixed by finding a more suitable optimisation algorithm, initial parameter values, or data transformation. These options are available within our web tool LAG PHASE DURATION CALCULATOR. However, we highlight the fact that the lag phase duration estimation is dependent not only on the selected method but also on multiple parameters of that method which tend to be underreported within experimental studies.

## CONCLUSIONS

The results presented here emphasize that while determining lag phase duration, all steps (pre-processing, choice of a method, and parameters) influence the final outcome. We would like to highlight that all these steps should be thoughtfully reported to ensure data reproducibility and credibility. Even the simplest tangent method requires specifying some details such as (i) if the initial biomass is represented by the first measurement or the minimal value, or (ii) the number of data points (or time frame) taken to draw the tangent line - whether a single point (e.g. [23]) or multiple points from the exponential phase (e.g. [7]) were used.

Within this publication, we came out with two solutions to facilitate the process of reproducible lag phase duration determination. First, we designed a decision tree (Fig. 7), which helps to choose the method best suited to one’s experimental conditions, taking into account various technical limitations and data imperfections. Second, we have developed an online tool, which will help to directly compare the lag phase duration estimated by different algorithms (i.e. combinations of methods, parameters, and data pre-processing techniques). The tool allows parameter adjustments and data pre-processing. Moreover, we share our code on GitHub [https://github.com/bognabognabogna/microbial_lag_calulator] so that it can be further developed and customised. In particular, we share the function “Calculate.Lag” which can be taken directly to the R environment. We perceive our tool as an initial point to further improvements made by the scientific community so that any new potential challenges can be solved in a reproducible way.

## METHODS

### Experimental growth curves

The empirical growth curve data used here were a part of a project described in Opalek *et al*. (2022) [19]. All growth curves were acquired by growing *Saccharomyces cerevisiae* S288C cultures in rich YPD media at 30°C. The populations were starved in various conditions before the growth procedure (see details in Supplement). The populations’ densities were monitored by absorbance measurements (optical density (OD), 600 nm in SpectraMax iD3) taken every half an hour. Then, the biomass was calculated according to the equations:

For OD measurements below 0.1: 10^(1.885845 + 28.72096×OD)^

For OD measurement equal or above 0.1: 3566518×OD + 26754296×OD^2^ -19881567×OD^3^

### SIMULATED GROWTH CURVES

Growth curves shown in Fig. 1 were simulated from the models described in the Supplement, and for time points spanning every 6 minutes between 0 and 24 hours. The parameters used are listed in the Supplementary Table 3.

### LAG DURATION CALCULATION METHODS

Let *t* denote time, *λ* denote lag duration, B denote biomass, and let *t*_*i*_ denote the i’th time point of the growth curve. If we define *N* = ln(*B*), then 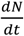 can be understood as the growth rate.

Additionally let *B*_0_ denote the first observation of biomass, and *N*_0_ = ln(*B*_0_)

#### Tangent to max. growth line

The derivative of *lN* is approximated using to the central scheme i.e.

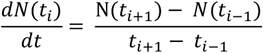

Then the maximal growth rate is defined as the maximum value of 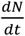 found within the growth curve i.e.: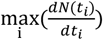. The first point at which growth rate is maximal is then denoted as (*t*_max.*growt*h_,*N*^max.*grwoth*^). Finally, a tangent line to this point is calculated.

Subsequently, lag duration is defined as the time when that tangent line crosses the *N*_0_ line. This value can be calculated as:

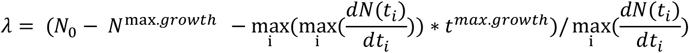

#### Tangent to max. growth point

The first point at which growth rate is maximal is calculated as above and it is denoted as (*t*^max,*growth*^, *N*^max,*grwoth*^). Then *n* points around that point are taken and a regression line is calculated using the *lm* function from the *stats* package (R). Subsequently, lag duration is defined as the time when that regression line crosses the *N*_0_ line. Within the manuscript n = 3. However, within our web tool this value can be modified by the user. Moreover, the value of *N*_0_ can be defined either as the first or the minimal log(biomass) value.

#### Parameter fitting to logistic model

All parameters of the logistic model (described in Supplement) are fitted simultaneously to the growth curve data using the *nlsLM* function from *minpack*.*lm* package (R), and the Levenberg-Marquardt algorithm, max.iter = 100. The initial values are estimated from the data and the following initial values: initial K = max(*B*), and initial lag = lag as calculated with the tangent method, and the initial growth rate is fitted to the data points that are likely to be in exponential growth i.e. those where biomass is between *B*_0_ + 0.2(max(*B*) − *B*_0_) and *B*_0_ + 0.8(max(*B*) − *B*_0_). All these initial values can be adjusted within the web tool.

#### Parameter fitting to Baranyi model

All parameters of the Baranyi and Roberts model (described in Appendix) are fitted simultaneously to the growth curve data using the *nls* function from *stats* package (R) and the *baranyi* formula from the *nlsMicrobio* package. We use the Gauss-Newton algorithm, max.iter = 100, and the following initial values: init.LOG10Nmax = max(log_10_(*B*), init.LOG10N0 = log_10_(*B*_0_), and the initial growth rate is fitted to the data points that are likely to be in exponential growth i.e. those where biomass is between *B*_0_ + 0.2(max(*B*) − *B*_0_) and *B*_0_ + 0.8(max(*B*) − *B*_0_). All these values can be adjusted within the web tool.

#### Max. growth acceleration

The second derivative of *N* is calculated approximated with the central scheme i.e.

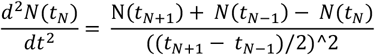

Then the max. growth acceleration point is defined as the maximum value of 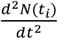 found within the growth curve i.e.: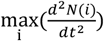. Consequently, lag duration *λ* is defined as the minimum time *t*_M_ at which 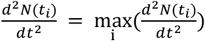.

#### Biomass increase

Let us define:

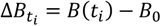

Lag duration *λ* is defined as the minimum time *t*_*M*_ at which 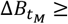 threshold

Finally, if the lag calculated by any of the method above turn out negative (which indeed may happen in the tangent and parameter fitting methods) we assume it is equal to 0.

Throughout this manuscript threshold = 10^5^. However, within our web tool this value can be modified by the user. Moreover, the value of *B*_0_ can be defined either as the first or the minimal biomass value.

Additionally in case the lag estimate is negative, NA is returned.

#### The code used to calculate the lag according to various methods can be found on GitHub https://github.com/bognabognabogna/microbial_lag_calulator

## ACKNOWLEDGMENTS

We would like to thank Wolfram Moebius, Ryszard Korona, Joanna Rutkowska, Aleksandra Walczak, Hanna Tutaj and Adrian Piróg for the discussion and their valuable comments. The research was funded by the Priority Research Area BioS under the program Excellence Initiative – Research University at Jagiellonian University in Krakow to BJS; by the Polish National Agency of Academic Exchange, grant number PPN/PPO/2018/00021/U/00001 to BJS, the programme “Excellence Initiative–Research University” at the Jagiellonian University in Kraków, Poland (grant number U1U/W18/NO/28.07) to MO; the National Science Centre, Poland, the OPUS grant to D.W.-S. (grant number 2017/25/B/NZ8/01035); the Biology Department research subsidies (grant number N18/DBS/000019 to MO and DWS).

## AUTHOR CONTRIBUTION

Conceptualization M.O., B.J.S., D.W.-S.; Empirical data M.O.; Data visualization B.J.S., M.O.; B.J.S. developed the web server MICROBIAL LAG PHASE DURATION CALCULATOR; Writing-original draft preparation - M.O, B.J.S.; Writing - reviewing B.J.S., D.W.-S.

All authors have read and agreed to the published version of the manuscript

## SUPPLEMENT

### Formulations of mathematical models used to simulate the growth curve data

1. Exponential model

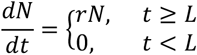

which can be solved as

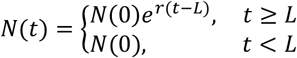

where N(t) is the population size at time t, L is the lag phase duration, and r is the growth rate (constant in time).
2. Logistic model

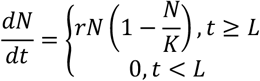

where N(t) is the population size at time *t, L* is the lag phase duration, *r* is the growth rate (constant in time), and *K* is the saturation constant (i.e. the maximum population size).
3. Monod Model

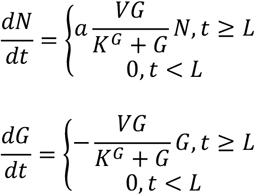

where N(t) is the population size at time t, G(t) is the limiting nutrient (for example glucose) concentration at time *t, L* is the lag phase duration. V denotes the maximal rate of the glucose uptake pathway, and *K*^*G*^ denotes the Michaelis-Menten constant (so that if 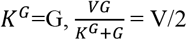). Moreover, the efficiency of converting glucose into biomass is described by a parameter a, which for simplicity is assumed to be constant.
4. Baranyi and Roberts Model

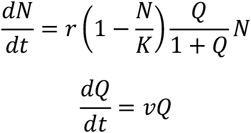

Where N(t) is the population size at time *t, K* is the saturation constant (i.e. the maximum population size), and *Q* represents the physiological state proportional to the concentration of some critical substrate.

## The growth curve data

Supplementary Table 1 gathers information about the populations used as examples (Fig. 2, Fig. 3) for empirical growth curves within this publication (Fig. 2, 6). We choose growth curves that demonstrate possible challenges in analysis, such as: biomass drop at the beginning of measurements (curve_1, curve_6), prolonged lag phase (curve_7, curve_8, curve_11), lack of easily-visible lag phase (curve_3), slow biomass increase (curve_8, curve_9), lack of exponential growth phase (curve_11), turbulent measurements in biomass (curve_10).

The data was originally collected to analyse possible advantages of phenotypic heterogeneity within a starving clonal population and it was used in Opalek *et al*. 2022. In particular, the growth curves come from three artificially prepared *S*.*cerevisiae* population types: Q monoculture, NQ monoculture, and mix culture. The Q (growth-arrested quiescent cells) and NQ (non-quiescent) cells are distinct phenotypes that can be separated from the stationary phase population (Allen *et al*. 2006). The populations had experienced starvation in complex (spent rich YPD medium) or simple (H_2_0) environments and had been regrown in rich media afterward for fitness analysis. For details, see Opalek *et al*. 2022.

The data was not analysed nor visualised in the form presented in this publication.

**Supplementary Table 1.** The summary of starvation conditions, chronological age, and phenotypic states of populations used to acquire growth curves analysed within this publication.

| CURVE ID | STARVATION MEDIA | STARVATION DURATION | CELL TYPE |
| --- | --- | --- | --- |
| curve_1 | complex environment | 7 days | NQ monoculture |
| curve_2 | complex environment | 7 days | NQ monoculture |
| curve_3 | complex environment | 7 days | natural mix culture |
| curve_4 | simple environment | 14 days | NQ monoculture |
| curve_5 | complex environment | 14 days | Q monoculture |
| curve_6 | complex environment | 14 days | Q monoculture |
| curve_7 | simple environment | 35 days | NQ monoculture |
| curve_8 | complex environment | 35 days | NQ monoculture |
| curve_9 | complex environment | 42 days | NQ monoculture |
| curve_10 | simple environment | 42 days | Q monoculture |
| curve_11 | simple environment | 42 days | NQ monoculture |

**Supplementary Table 2.**
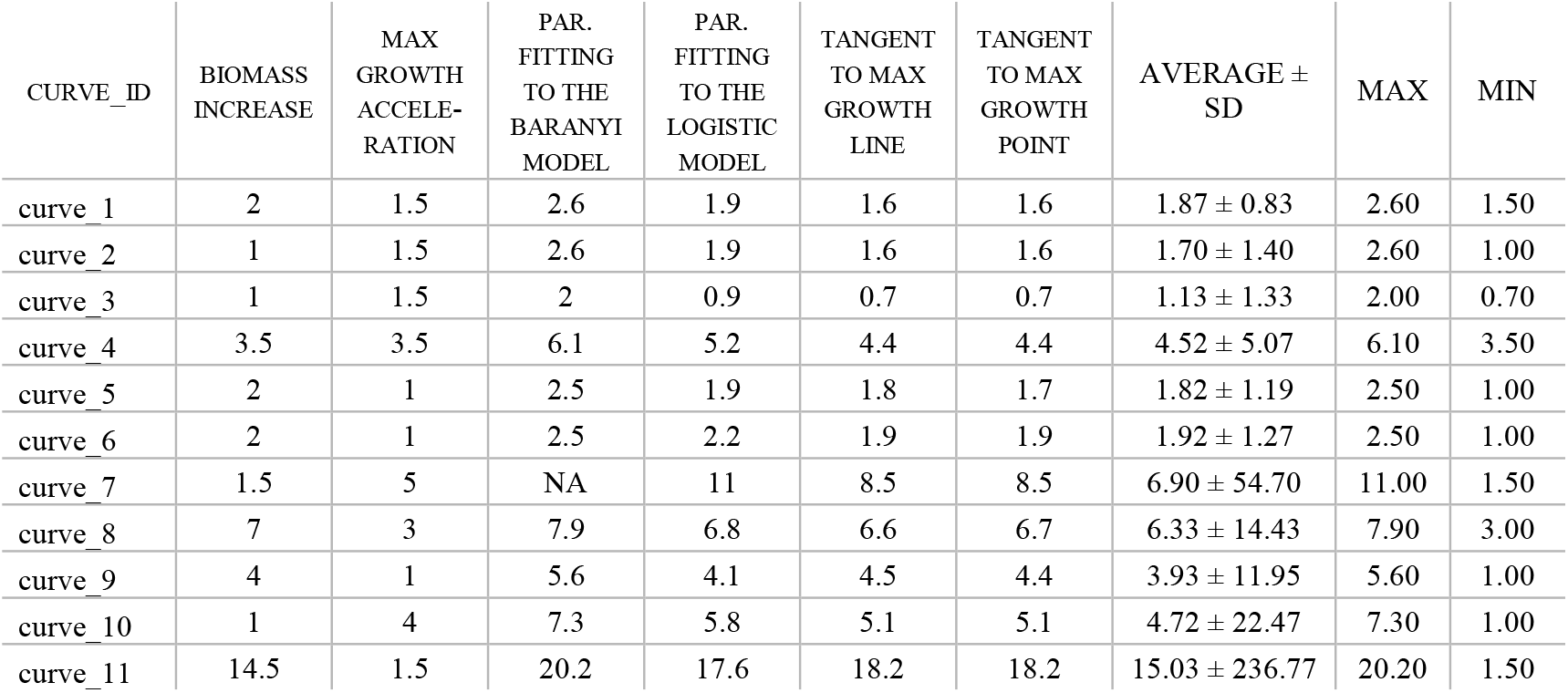
The summary of lag phase durations calculated by various methods (columns). The table compares the similarities of results for each of the curves. The numbers are given in hours.

**Supplementary Table 3.** Default parameters used to simulate data from models described in section Formulations of mathematical models used to simulate the growth curve data.

| Parameter name | Default value |
| --- | --- |
| Growth rate $r$ (exponential and Baranyi model) | $0.1 [h^{-1}]$ |
| Max. growth rate $r$ (logistic model) | $0.25 [h^{-1}]$ |
| Carrying capacity $K$ (logistic model) | $5 * 10^6$ [cells/mL] |
| Initial biomass $N_0$ | $10^6$ [cells/mL] |
| Lag $L$ in logistic, exponential, and Monod model | 2.5 [h] |
| Initial physiological state in Baranyi model $Q_0$ | 3.52 [units] |
| Efficiency of substrate uptake in Monod model $a$ | $3*10^5$ [cells/mmol substrate] |
| Maximal substrate uptake rate in Monod model $V$ | $1.5*10^{(-5)}$ [mmol substrate / cells × h] |
| Michaelis-Menten constant in Monod model $K^G$ | 300 [mmol substrate] |

**Supplementary Figure 1.**
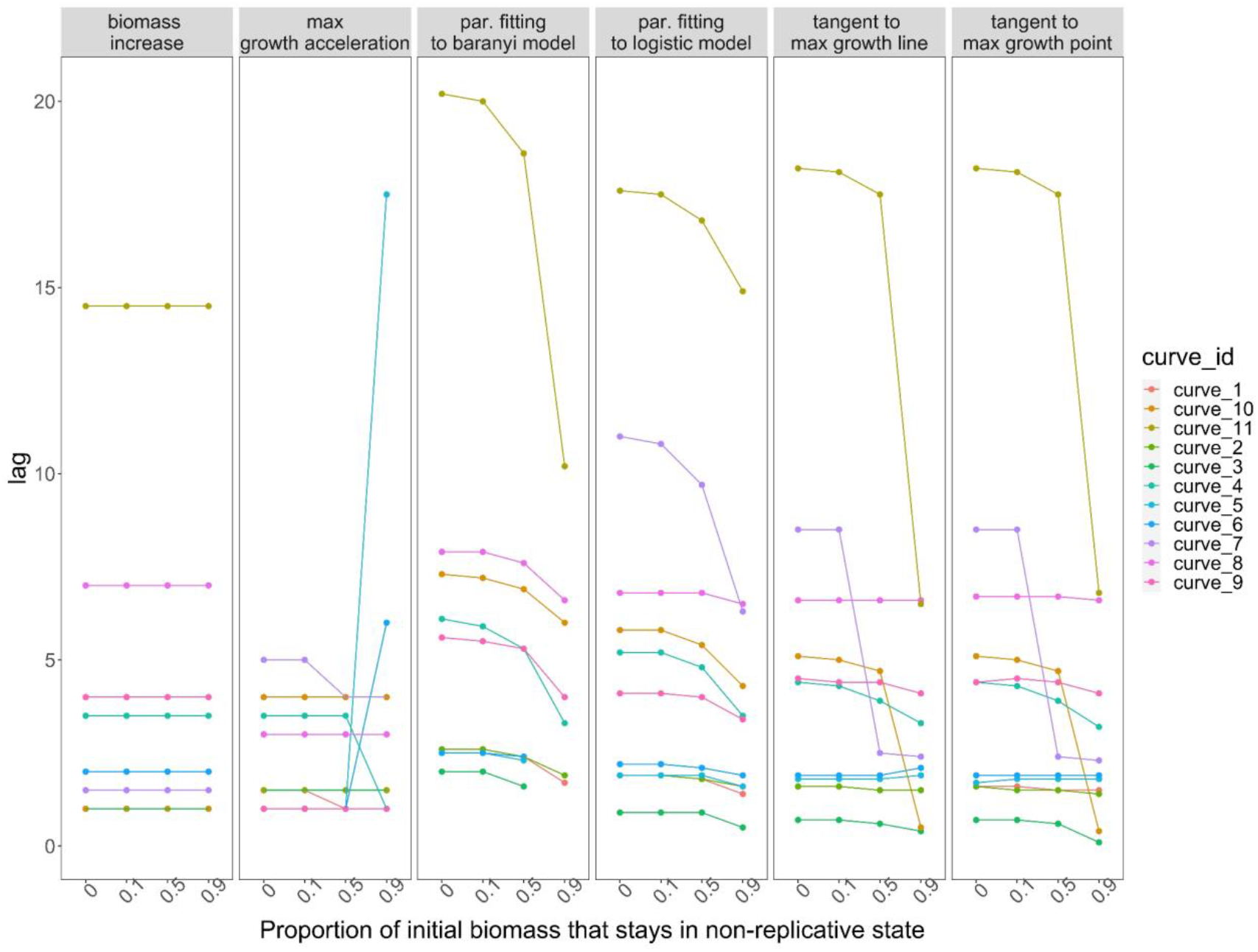
Lag duration as calculated for 11 experimental growth curves with various methods, assuming that a percentage of the initial population is non-replicative.

In case of growth curves, where there is no biomass increase for prolonged time (e.g. curve_11), the proportion of alive vs senescent (permanently non-replicative cells) highly impact estimated lag duration. If all cells are capable of proliferating (x = 0), then the observed lack of biomass increase is indeed a lag phase, however if a substantial fraction of population is unable to proliferate (x = 0.9), then the remaining 10% of alive cells start proliferate earlier (lag = 5-10 h, depending on the method (columns)), however the increase in biomass is small to be marked as the lag phase end. That is why the proportion of dead cells should be also monitored when analysing growth curves.

**Supplementary Figure 2.**
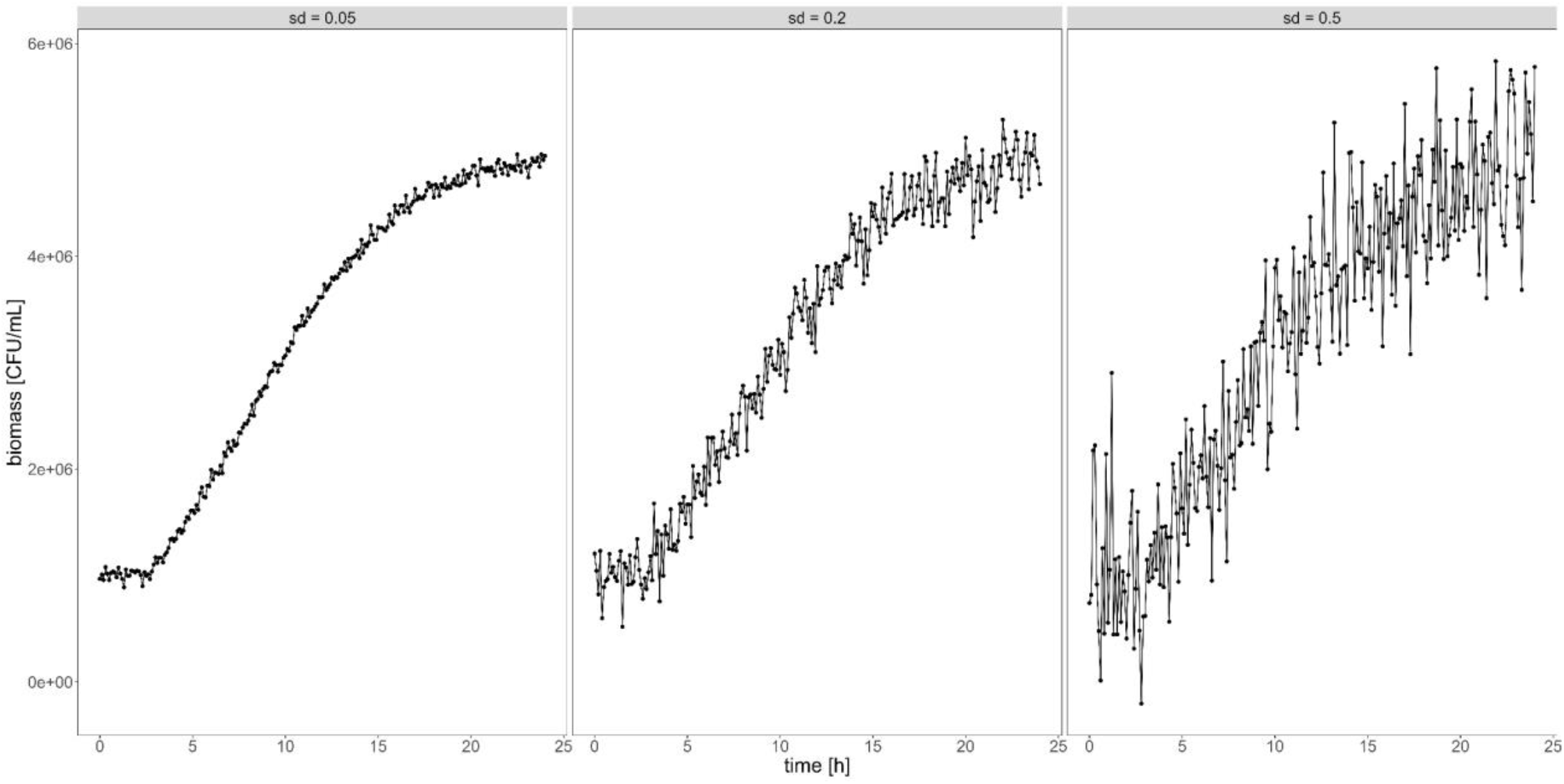
The visualization of the impact of introduced noisiness (sd) on growth curve shape. The noise was simulated from the random distribution with mean = 0 and standard deviation standardized by the initial biomass B0

**Supplementary Fig. 3.**
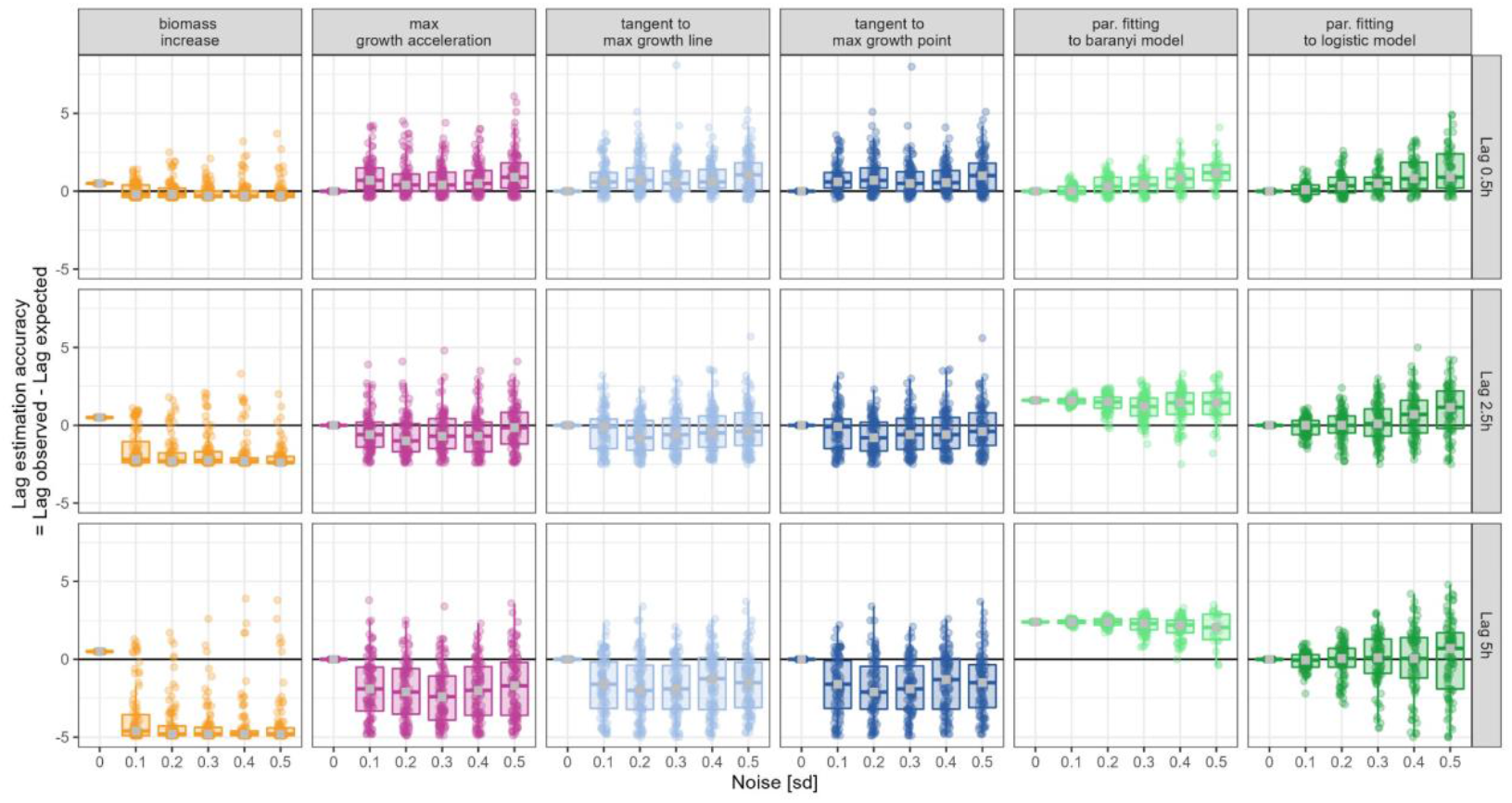
Impact of real lag phase duration on calculation accuracy. For each sd (noisiness, x-axis), expected lag length (rows), and lag calculation method (columns), 100 simulations were conducted (points). For each row, the true (expected) lag was set as a different value, e.g. in the first row, the true lag was set as 0.5h, and the y = 0 correspond to lag = 0.5h, while in the second row, y = 0 correspond to lag = 2.5h. Boxplots illustrate the distribution of points, and the median value (grey square). All points located above y = 0, show these calculations where lag phase length was overestimated, similarly, all points below y = 0 show calculations where lag duration was underestimated.

Parameter fitting to Baranyi model systematically overestimates lag phase duration. For short lags (first row), all but biomass increase methods tend to overestimate lag duration when noisiness increase, while for long lags (last row), all but parameter fitting to logistic model methods underestimate lag duration when noisiness increase. Median value (grey square) for the parameter fitting to the logistic model is closest to y = 0 what indicates that this method is least biased.

**Supplementary Fig. 4.**
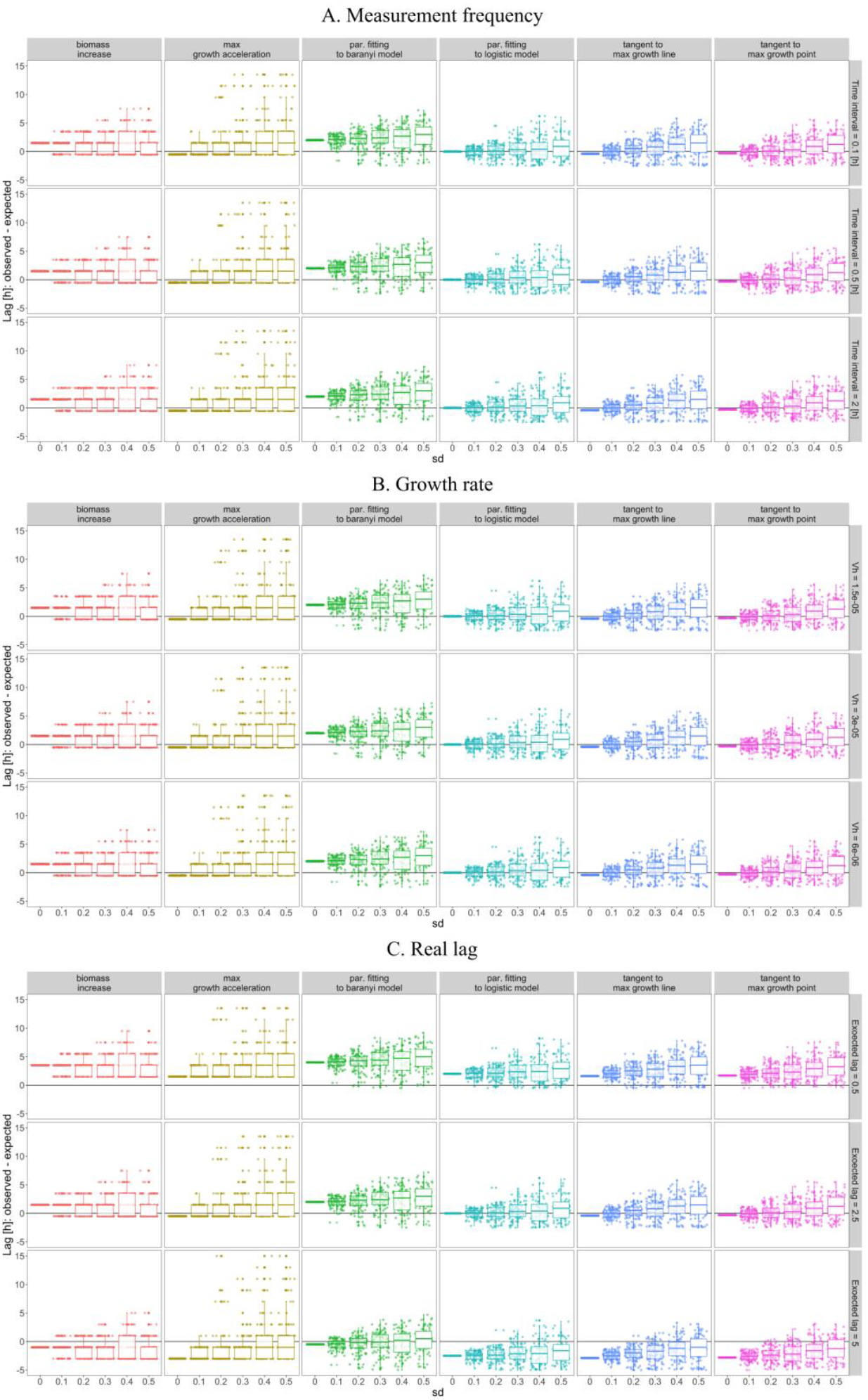
Testing robustness of results for data simulated via Monod model. The biases and accuracies of the methods show the same pattern as described in the manuscript (Fig. 3, Fig. 4, Supp. Fig. 3), and as such, the outcomes are not driven by the model itself.

**Supplementary Fig. 5.**
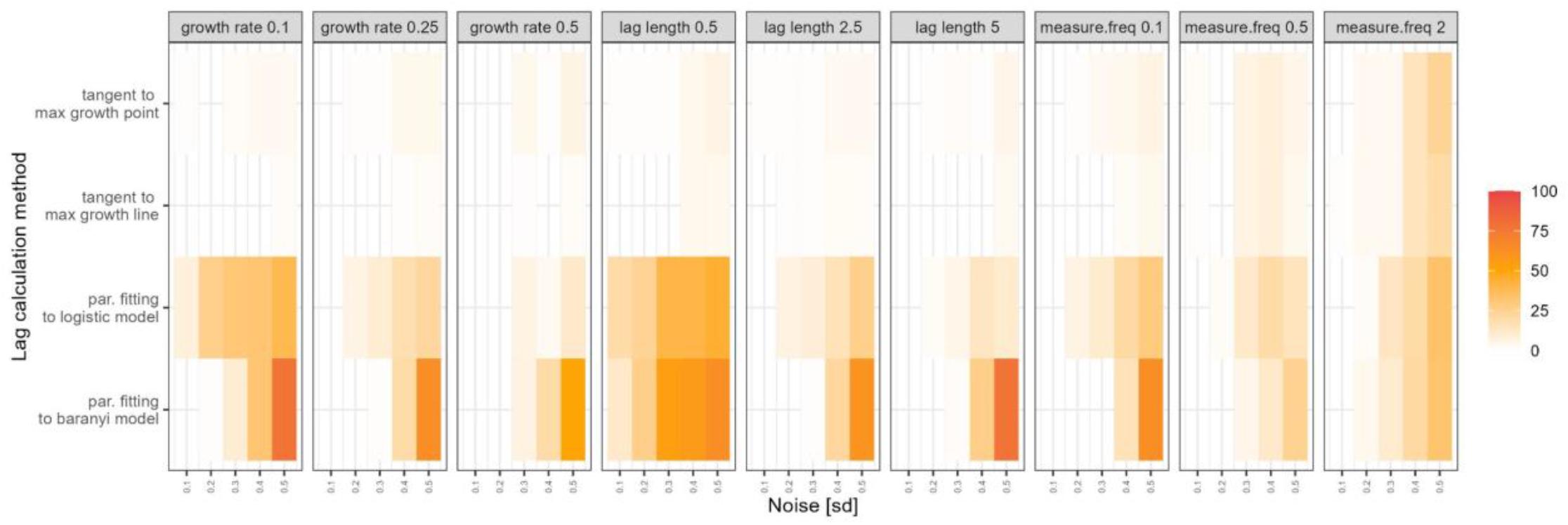
Frequency of NA value generated (indicating failure of the lag estimation) out of 100 simulated growth curves (the curves described in section: Testing the sensitivity of lag determination methods to data noisiness). In case of parameter fitting to model NA means that the fitting didn’t converge to any solution, while for tangent method NA is generated if lag was estimated as a negative value. No filling indicates that all 100 lag estimations values were generated. For biomass increase and max growth acceleration methods all lag estimations were successful and as such they were excluded from the graph. The maximum of 78 failed lag estimations were generated twice for parameter fitting to Baranyi model: for data simulated with slow growth rate (0.1) and long lag phase (5h), both with and high noise (sd = 0.5).

